# Ad-Seq, a genome-wide DNA-adduct profiling assay

**DOI:** 10.1101/364794

**Authors:** Olivier Harismendy, Stephen B. Howell

## Abstract

**Summary:** Carcinogens form adducts with the DNA which, when not properly repaired, can lead to mutations and drive oncogenesis. The identity, sequence specificity and mutagenicity of most DNA-adducts is however poorly understood and current molecular assays are limited in their scope and scalability. We present a novel genome-wide DNA adduct sequencing (Ad-Seq) assay to map the location of DNA-adducts at single-nucleotide resolution. Ad-Seq enriches for DNA fragments containing nuclease digestion resistant DNA-adducts. The genomic location of the resulting reads is aggregated in a quantitative profile showing the DNA-adduct sequence context. Ad-Seq is quantitative and confirms known specificity of damages from Ultra-Violet light (di-pyrimidine) and cisplatin (AG and GG di-purines). Furthermore, in cells, Ad-Seq profile can be compared to chromatin segments to show that cisplatin associated adducts are depleted in open and active chromatin regions. The Ad-Seq assay can therefore generate a broad DNA signature of DNA damage and, by comparing to mutagen exposure or downstream mutational profile and signatures, be used to improve our understanding of cancer molecular etiology.

---

The formation and repair of DNA-adducts is the earliest process in oncogenesis following exposure to most mutagens[1]. Unsuccessful or inaccurate repair leads to somatic mutations, large chromosomal aberrations and/or apoptosis. DNA adduct formation is the main mechanism of action of a number of genotoxic chemotherapies such as platinating or alkylating agents. In the absence of apoptosis, the mutations rising after inaccurate DNA repair can drive cellular proliferation and cancer progression. The resulting tumors often display specific mutational signatures characterized by the type and sequence context of the nucleotide substitutions observed and associated with putative mutagenic stress and processes[2]. The mutagen-associated mutational signatures are likely the final result following mutagen activation, reaction with the DNA, faulty repair of the damaged nucleotide and mutation propagation through cellular proliferation. Assays to quantify DNA-adducts, such as P^32^ post-labelling or Liquid Chromatography Tandem Mass Spectrometry, do not provide information on the nucleotide context or are restricted to a narrow spectrum of defined chemical structures[3]. Recent DNA sequencing-based assays can locate specific sites of damage to the DNA and have been applied to map damage from ultra-violet (UV) light or a restricted set of chemical mutagens (Table S1 and associated references). Some of these methods have a low resolution (DNA interval) and all require either immunoprecipitation of the DNA-adduct, when a specific antibody is available, or an adduct-specific chemical treatment to allow the isolation and sequencing of the damaged DNA fragment.

We developed the DNA-adduct sequencing assay (Ad-Seq) to overcome these limitations and allow the unbiased mapping of DNA-adducts at single nucleotide resolution (Figure 1a). Ad-Seq enriches a genome-wide library for DNA fragments containing intra-strand nucleotide dimer adducts. The digestion of a mechanically fragmented genome-wide DNA library using Lambda and Rec-Jf exonucleases generates a library of single-strand DNA enriched for fragments resistant to the digestion due to the presence of an adduct at their 5’ end. This digestion is followed by the 3’ C-tailing of the fragment, the linear amplification of the reverse strand from a HG9 primer up to a position preceding the adduct and the ligation of the 2N adapter containing a random two-nucleotide overhang. Both primer and adapter contain sequences required for the subsequent amplification and sequencing by synthesis (e.g. Illumina chemistry).

**Figure 1:**
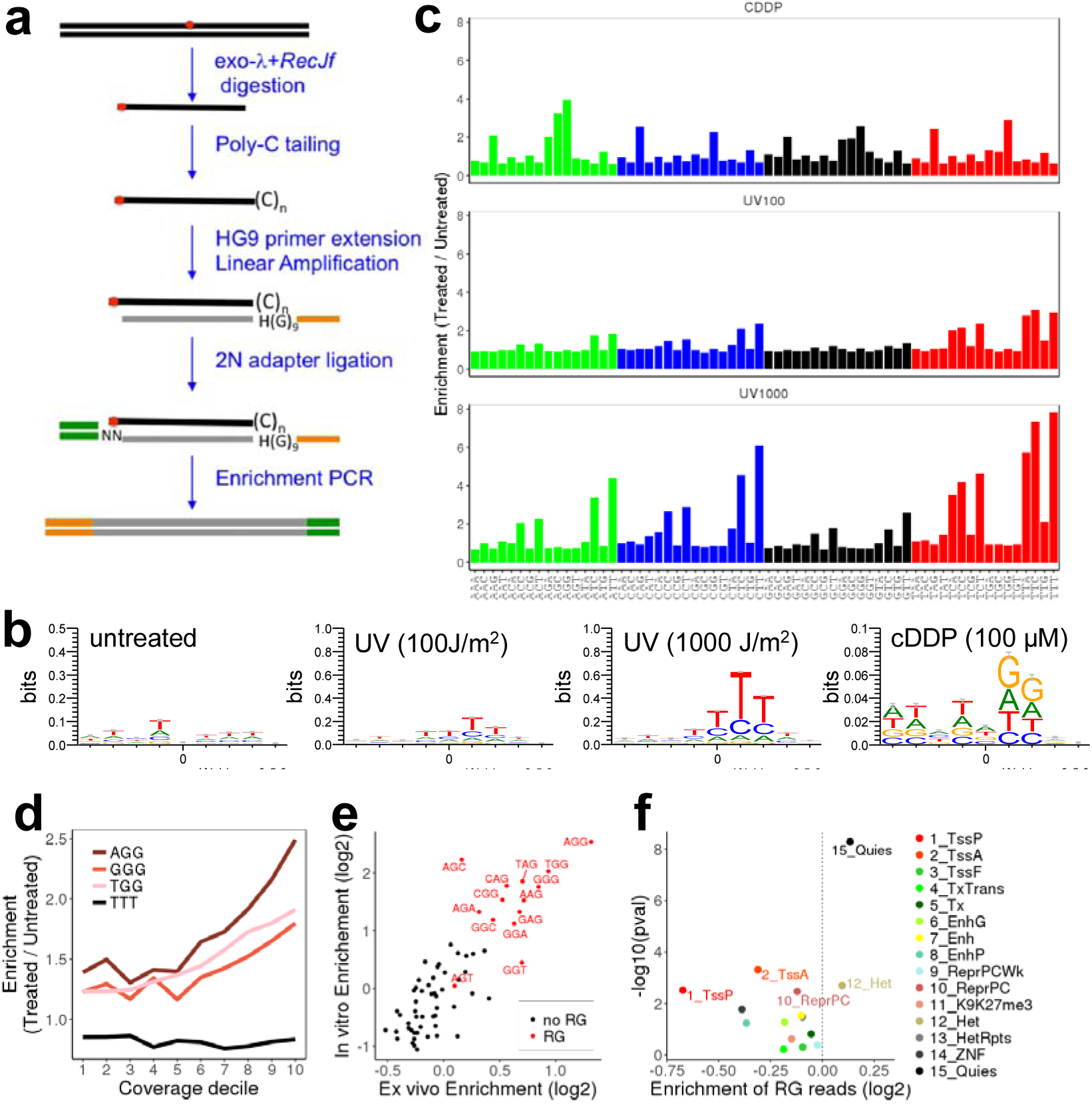
Ad-Seq, a genome-wide DNA-adduct profiling assay. **(a)** Summary of Ad-Seq experimental workflow. The intra-strand DNA adduct is indicated in red. **(b)** DNA sequence logo (9-mer) at genomic positions the most covered (top decile) by Ad-Seq reads (read start at x=0). **(c)** Enrichment of trinucleotide patterns at the start of the mapped sequencing reads in the *in vitro* treated compared to untreated DNA. **(d)** Enrichment of three types of dinucleotides at the read start of the *ex vivo* cDDP-treated DNA as a function of the coverage depth (x axis). **(e)** Enrichment at all trinucleotide pattern compared between ex vivo (x-axis) and in vitro (y-axis) cDDP treatment. Enrichment was calculated using condition-matched untreated controls using reads from the top decile of coverage depth. **(f)** Enrichment of cDDP adducts (top 10% most covered genomic position with RG dimer in the starting trinucleotides) in 15 different types of chromatin regions of the IMR90 cell line relative to an untreated sample. Significant types (p<0.01) are labelled. Abbreviations available at the Epigenome Roadmap UCSC Hub [9].

We first carried out Ad-Seq on naked DNA treated *in vitro* with UV light (100 and 1000 J/m^2^) or cisplatin (cDDP: 1 μM for 4 h). The sequence observed at the 5’ end of the reads mapped to the genome was consistent with the known chemistry of the mutagen (Figure 1b): UV induces the formation of pyrimidine dimers[4], reflected by an enrichment of C and T at the beginning of Ad-Seq reads. Similarly, cDDP creates mainly intra-strand adducts at AG and GG (referred to as RG) di-puric sites[5], consistent with A and G enrichment observed at the beginning of Ad-Seq reads, extending into the third nucleotide of the read. Consistently, trinucleotide sequences at the start of the reads are enriched for the expected pattern compared to the untreated DNA (Figure 1c), with pyrimidine dimers (YY) significantly enriched in UV treated DNA and RG purine dimers in cDDP treated DNA. The enrichment of YY is stronger for high UV exposure (6.9 vs 2.6-fold for TT) suggesting that the Ad-Seq profile is quantitative. Interestingly, while UV-induced mutations consist mostly of CC to TT substitutions[6], the UV Ad-Seq profile is enriched at all types of di-pyrimidines. The difference is likely the consequence of a more error prone repair of cyclobutane pyrimidine dimer occurring at CC dimers and more efficient translesion synthesis at TT [6,7]. This observation highlights the important information on mutational processes that can be gained by comparing Ad-Seq and mutational profiles.

We next treated a lung fibroblast cell line (IMR90) *ex vivo* with 100 μM cDDP for 4 h before DNA extraction and Ad-Seq profiling (Figure 1a). The enrichment of trinucleotide sequence patterns containing RG di-purines increased with coverage depth (Figure 1d) and correlated with the one observed *in vitro* (Figure 1e). We evaluated the distribution of reads across different functional segments of the chromatin previously established in the same cell line [8,9]. Compared to untreated cells, reads with RG dimers in their starting tri-nucleotide sequence were significantly depleted at active and poised transcription start sites (p<0.005) and enriched in the quiescent chromatin (p<10^−8^, Figure 1f). This result is consistent with a more active repair observed in active and open chromatin regions [10].

Ad-Seq represents a novel molecular assay to discover and characterize DNA signatures associated with mutagenic exposures. In contrast to single agent / single adduct approaches, Ad-Seq profiling represents promising approach to reveal associations between mutagen exposure and their biological and genetic consequences and exploring how cellular processes mediate the genesis of somatic mutations or modulate the activity of genotoxic chemotherapies.

## Acknowledgments

We are grateful to Brian Woo, Christopher Fang and the Institute for Genomics Medicine Genomics Center for their technical assistance. This work was supported by a Pilot award from the San Diego Center for Systems Biology (NIH award P50GM085764) and by the NCI IMAT award R21CA177519.

